# Noise Reconstruction & Removal Network: A New Architecture to Denoise FIB-SEM Images

**DOI:** 10.1101/2021.05.27.446051

**Authors:** Katya Giannios, Abhishek Chaurasia, Guillaume Thibault, Jessica L. Riesterer, Erin S. Stempinski, Terence P. Lo, Bambi DeLaRosa, Joe W. Gray

## Abstract

Recent advances in Focused Ion Beam-Scanning Electron Microscopy (FIB-SEM) allows the imaging and analysis of cellular ultrastructure at nanoscale resolution, but the collection of labels and/or noise-free data sets has several challenges, often immutable. Reasons range from time consuming manual annotations, requiring highly trained specialists, to introducing imaging artifacts from the prolonged scanning during acquisition. We propose a fully unsupervised Noise Reconstruction and Removal Network for denoising scanning electron microscopy images.

The architecture, inspired by gated recurrent units, reconstructs and removes the noise by synthesizing the sequential data. At the same time the fully unsupervised training guides the network in distinguishing true signal from noise and gives comparable results to supervised architectures. We demonstrate that this new network specialized on 3D electron microscopy data sets, achieves comparable and even better results than supervised networks.

## 1 Introduction

Recent advances in Focused Ion Beam-Scanning Electron Microscopy (FIB-SEM) have led to unprecedented biological tissue visualization and analysis, as well as understanding of cellular ultrastructure and cell-to-cell interactions (Xu, Hayworth, Lu, Grob, Hassan, Garcia-Cerdan, Niyogi, Nogales, Weinberg & Hess 2017). High-resolution FIB-SEM data sets often consist of volumes sliced into thousands of 6K × 4K images with 4nm resolution per voxel, allowing a 3D reconstruction of a fraction of tissue volume. Among others, one of the problems faced during analysis of such data is the presence of noise. Depending on the tissue type, sample preparation, acquisition settings, detector used, etc., the images may contain a significant quantity of noise making any further analysis tedious or even impossible (Kubota, Sohn & Kawaguchi 2018, Liu, Sun, Gao & Li 2018).

By definition, image denoising is the process of taking a noisy image *x* and separating the noise *n* from the true signal *s*: *x* = *s* + *n*. Following the typical assumption for the noise (Foi, Trimeche, Katkovnik & Egiazarian 2008, Wu, Gong, Kim & Li 2019) and taking the microscope’s characteristics into account, we can assume that the noise in microscopy images is: (i) a zero-mean random noise, that is for any pixel the noise is a discrete random number added to the pixel ‘true value’; (ii) each pixel noise is independent, so the noise value at any pixel does not depend on the noise at any other pixel, but it is signal dependent. While the noise is random and independent, the signal is not and this is what, typically, denoising methods rely on.

In recent years, deep learning methods have established themselves as powerful analytical tools in machine learning (Arik, Chrzanowski, Coates, Diamos, Gibiansky, Kang, Li, Miller, Ng, Raiman et al. 2017, Jing, Yang, Feng, Ye, Yu & Song 2019, Karpathy & Fei-Fei 2015, Oord, Dieleman, Zen, Simonyan, Vinyals, Graves, Kalchbrenner, Senior & Kavukcuoglu 2016). Specifically, Convolutional Neural Networks (CNNs) have been used for various tasks including detection (Lin, Goyal, Girshick, He & Dollár 2018, Liu, Anguelov, Erhan, Szegedy, Reed, Fu & Berg 2016, Redmon & Farhadi 2018, Ren, He, Girshick & Sun 2016, Tan, Pang & Le 2020), segmentation (Chen, Papandreou, Schroff & Adam 2017, Chen, Zhu, Papandreou, Schroff & Adam 2018, Ronneberger, Fischer & Brox 2015, Tao, Sapra & Catanzaro 2020, Yuan, Chen & Wang 2019), super resolution (Sun & Chen 2020), etc. In the field of denoising, CNNs have been very useful (Kim, Chung & Jung 2019, Liu, Wu, Wang, Xu, Zhou, Huang, Wang, Cai, Ding, Fan & Wang 2019, Yu, Park & Jeong 2019) especially when the noise characteristics are unknown, making any mathematical modeling difficult. In this paper we apply a CNN technique to FIB-SEM acquired images, taking into account the relevant specifics.

FIB-SEM allows 3D imaging of biological fine structure at nanoscale resolution: a thin slice of the sample is removed with the ion beam and the newly exposed surface is imaged with the electron beam. That results in a sequence of images containing isotropic voxels down to 4nm. To improve the signal-to-noise ratio the researcher may decide to image the surface several times and utilize frame averaging.The downside of using that technique for noise reduction is the possibility of specimen charging and/or shrinking due to electron dose and the impact on the general acquisition time. An alternative will be using denoising techniques to improve the image quality. Traditionally, training neural networks for denoising demands pairs of noisy and clean images (ground truth images): (*x*_*j*_, *s*_*j*_). Theoretically, obtaining denoised images is possible by averaging multiple (up to hundreds) acquisitions of the same sample. As mentioned, this is not feasible with FIB-SEM. The challenges with obtaining ground truth images, as in many biological use cases, is motivation for developing and utilizing unsupervised techniques, such as a Noise2Noise approach (Wu, Gong, Kim & Li 2019, Lehtinen, Munkberg, Hasselgren, Laine, Karras, Aittala & Aila 2018).

We use a triplet of images (*x*_*j*−1_, *x*_*j*_, *x*_*j*+1_) as input in our proposed denoising process. The additional two images *x*_*j*−1_ and *x*_*j*+1_, that can be considered as technical replica of the image *x*_*j*_ because of the high resolution and the signal redundancy, are used to boost the signal *x*_*j*_ to denoise. We create two pairs (*x*_*j*−1_, *x*_*j*_) and (*x*_*j*+1_, *x*_*j*_) that are fed to the same network, which is then trained to map one noise realization to the other, using our modified Noise2Noise loss function. We refer to the proposed architecture as Noise Reconstruction and Removal Network (NRRN). The NRRN is applicable to the case of improving the image quality based on two or three scans of the same slice or denoising based on the two adjacent slices in the volume stack. Our major three contributions discussed in this paper are:

1. a novel noise reconstruction module with soft attention and signal boosting, that upon deployment on very large images (more than 24M pixels) homogeneously removes the noise,
2. a neural network architecture design using our noise reconstruction module with detailed performance analysis, and
3. updated noise2noise loss function (Wu, Gong, Kim & Li 2019) specifically designed for denoising FIB-SEM data.

In what follows we give a short overview of related works.Noise-to-clean (N2C) is the traditional supervised learning approach, where the models are trained with pairs of noisy and clean images (*x*_*i*_, *s*_*i*_) as inputs and targets respectively, where *x*_*i*_ = *s*_*i*_ + *n*_*i*_, so that the network learns to remove the noise. However, when the clean (ground truth) images are not available, the supervised N2C approach is not applicable. In 2018, Lehtinen et al. (Lehtinen, Munkberg, Hasselgren, Laine, Karras, Aittala & Aila 2018) introduced a new approach: the Noise2Noise (N2N). Instead of training a network to map noisy inputs to clean images, their N2N approach trains on pairs of independently degraded versions of the same training sample: 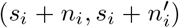. Authors demonstrate that predictions from networks trained with such N2N approach converge to the same predictions as traditionally N2C trained networks.

(Wu, Gong, Kim & Li 2019) further developed the N2N idea with a novel loss function applied to medical images with available pairs of noise realizations. The network was then trained to map one noise realization to the other, with a loss function that efficiently combined the outputs from both training subsets. A mathematical proof was provided to demonstrate that the proposed training scheme was equivalent to training with noisy and clean samples, empirically proving that the noise in the two subsets was random and spatially independent. Noise2Void (N2V)(Krull, Buchholz & Jug 2019) and the follow up Noise2Self (N2S) (Batson & Royer 2019) training schemes fill the gap when no clean target images or technical replica are available. If these architectures training are less restrictive, N2V authors mentioned that they do not outperform the N2N or N2C methods since they have less information available during training. In our case we take advantage of the volumetric structure of the data that lead to information redundancy, and we chose a N2N training style.

Independent of the training scheme, a lot of work has been done on the neural network architectures design. A very popular choice is the U-Net architecture (Ronneberger, Fischer & Brox 2015) and its variations. U-Net was first introduced for segmentation purposes and since has been widely extended in terms of applications as well as architectures. Alternatively, Remez et al. (Remez, Litany, Giryes & Bronstein 2017, 2018) used a fully-convolutional neural network architecture that exploits the gradual nature of the denoising process. Their DenoiseNet architecture calculates a “noise estimate” that is fine-tuned at each layer based on the previous layer output, and the results are then added to the input image. They have shown that the shallow layers handle local noise statistics, while deeper layers recover edges and enhance textures. Recurrent neural networks (RNN) have been widely developed and used in many applications, such as natural language processing and video analysis. RNNs handle temporal sequences of input, and perform the same processing at each element of the sequence while keeping memory of the previous computation.

Influenced by RNN principles, and more specifically its two most famous derivatives, GRU (Cho, van Merrienboer, Gulcehre, Bahdanau, Bougares, Schwenk & Bengio 2014, Yao, Cohn, Vylomova, Duh & Dyer 2015) and ConvLSTM (Shi, Chen, Wang, Yeung, Wong & Woo 2015, Yang, Xiong, Xu, Zhou, Xu, Chen, Park, Grbic, Tran, Chin, Metaxas & Comaniciu 2017), as well as DenoiseNet architecture, we propose a new denoising architecture that models the noise using information from two or three images. At each layer we accumulate the information from the additional images and calculate a noise component. During network training, we use an unsupervised noise2noise loss function adapted to the multi-image architecture - more details in subsection 2.1.2.

## 2 Results

### 2.1 Noise Reconstructing & Removal Network

The proposed Noise Reconstruction and Removal Network (NRRN) architecture and the training framework are designed to take advantage of the sequential nature of FIB-SEM data. NRRN uses specific units to fuse the additional information as well as fully unsupervised training to guide the network in distinguishing true signal from noise.

#### 2.1.1 Denoising module architecture

Typically, a FIB-SEM data set consists of *N* images representing *N* consecutive imaged slices (*x*_0_, …, *x*_*N*_). Our approach is to obtain a denoised image 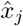 from three adjacent noisy images (or if available, three scans of the same slice). To do so, we consider the initial tissue sample into a data set of *N* − 2 independent triplets of images (*x*_*j*−1_, *x*_*j*_, *x*_*j*+1_). We split each triplet into two pairs (*x*_*j*−1_, *x*_*j*_) and (*x*_*j*_, *x*_*j*+1_) that are passed through two parallel identical branches of the proposed network NRRN, see Fig.1 A. For a pair of images (*x*_*j*_, *x*_*j±*1_), each branch (Fig. 1 B) consists of:

**Figure 1:**
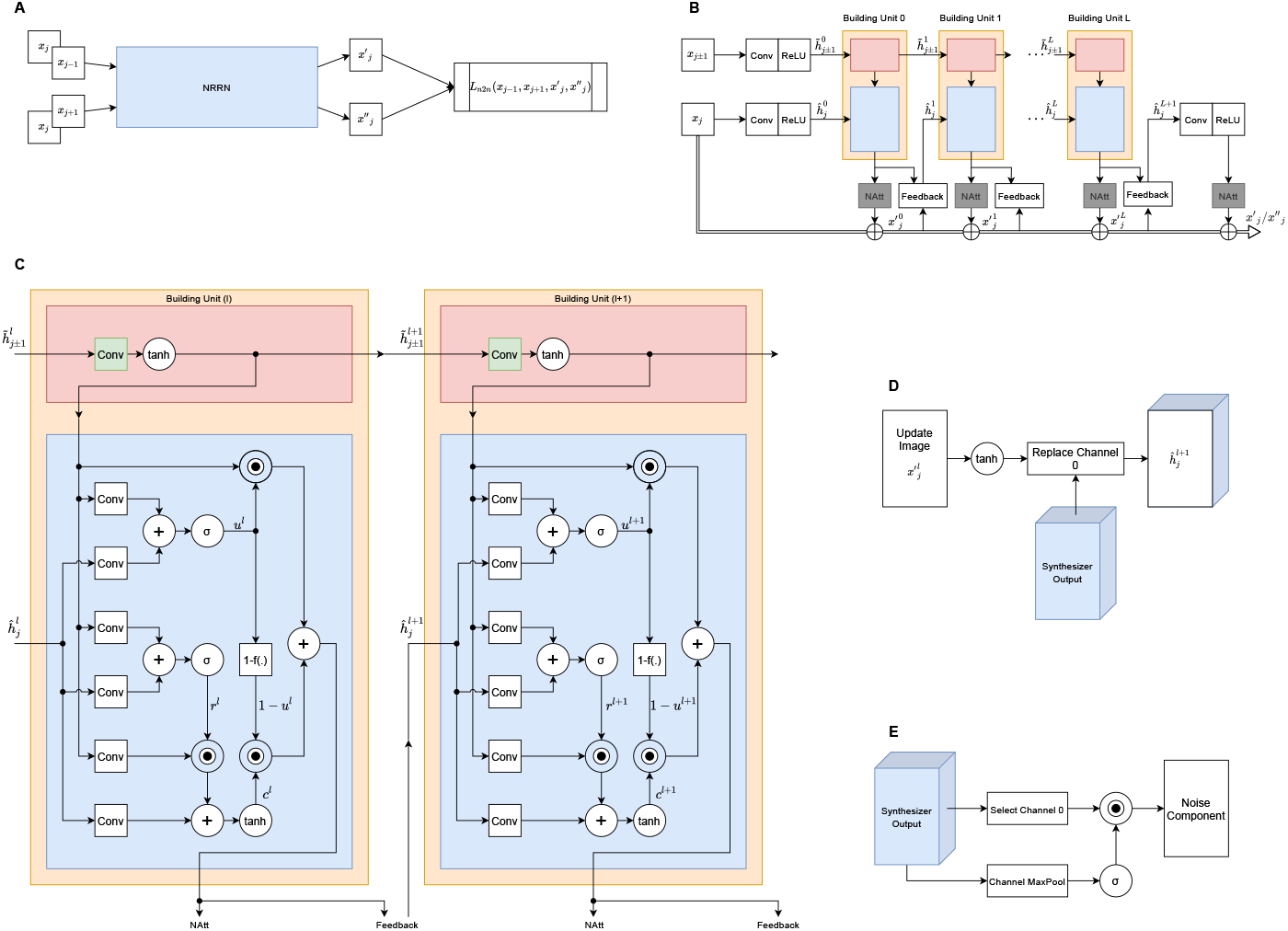
Noise Reconstructing & Removal Network Architecture. (A) Training Process; (B) Single Branch Architecture; (C) Building Unit; (D) The Feedback Block after the *l*^*th*^ BU; (E) The Noise Attention Block (NAtt)

- one convolution layer (3 × 3 kernel, 64 channels, and ReLU activation (see the Supplemental Material for definition)) applied to each of the two images
- *L* + 1 stacked layers made of a building unit (BU) coupled with noise attention (NAtt) and feedback blocks
- a final convolution layer (3 × 3 kernel, 64 channels, and ReLU activation).

By stacking BUs, the network architecture is an iterative denoising process comparable to DenoiseNet. Each BU models noise components (see Fig. 1 C) by accumulating noise characteristics from previous units and from updated input image after each stage. Within the BU *l*, the information flows through two major paths which have been highlighted by an upper and a lower block. While the path through the lower block 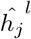 learns noise features from previous BU (*l* − 1) and cleaner input 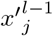, the upper block 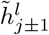 mostly learns information coming from the adjacent image *x*_*j±*1_ by performing bulk of operation and we call it synthesizer. Motivated by LSTM and GRU cells, but with a reduced computational overhead, a BU *l* contains two gates: an update gate *u*^*l*^ that decides how much of the previous/adjacent information (coming from 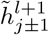) and the new one (1 − *u*^*l*^) needs to be kept; a reset gate *r*^*l*^ that regulates how much needs to be forgotten. This allows the synthesizer at every level to accept useful information coming from the *x*_*j±*1_ while simultaneously being aware about the updated image 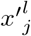.

After the synthesizer has learned the noise components, the new features are then passed to the NAtt block that filters the BU output and generates the noise component which is removed from the corresponding input image to give a new updated image 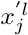 to the next BU. The key equations for every pair of images (*x*_*j±*1_, *x*_*j*_) are given in eq.1, where ◦ denotes the Hadamard product and

* denotes the the convolution operator:

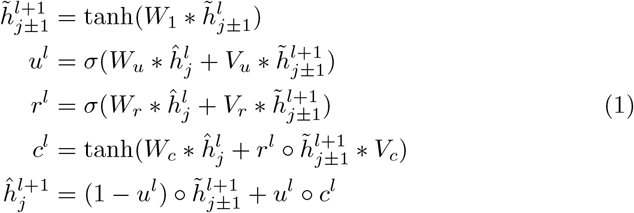

for *l* ∈ {0, …, *L*}, *j* − {1, …, *N* × *M*} and 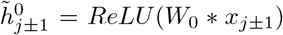 and 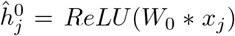 and *W*_1_ is with shared parameters across all BUs. The noise attention block (see Fig. 1E) receives a feature vector with 64 channels 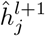 as input. It max pools through all channels to create an attention map that gets element-wise multiplied to the first channel of 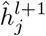. We observe that the output of the NAtt block represents component of the noise present in previous noisy image 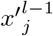.

Presumably, we have a less noisy image 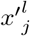 after each layer and we want to feed it back to the network. That is achieved in the feedback block (see Fig. 1 D) that works as a buffer by gathering information from both the updated image 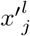 as well as the newly learned features 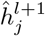 from the previous building unit. Thus the first channel of the synthesizer output 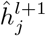 is updated by the Feedback block giving the input for next BU. For simplicity we use the same notation 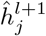, for both: synthesizer output and input of the next BU.

#### 2.1.2 Loss Function

In (Wu, Gong, Kim & Li 2019), showed that for a clean image *s*_*k*_ and two noisy realizations of it *x*_(*k*,1)_ = *s*_*k*_ + *n*_(*k*,1)_ and *x*_(*k*,2)_ = *s*_*k*_ + *n*_(*k*,2)_ the function

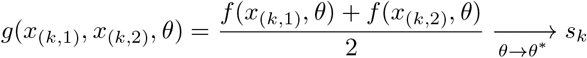

approaches the real signal *s*_*k*_ for, *θ*^*^ = *arg* min *L*_*n*2*n*_(*θ*) where

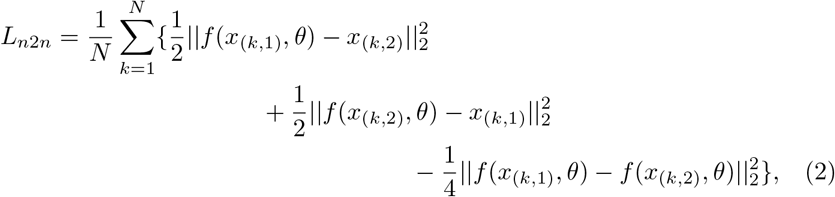

The only requirements for such loss function are that the noise *n*_(*k*,1)_ and *n*_(*k*,2)_ are with zero means and spatially independent.

In our case, we consider *N* − 2 triplets of images split into two pairs (*x*_*j*−1_, *x*_*j*_) and (*x*_*j*+1_, *x*_*j*_). The NRRN network maps each of these pairs to two outputs 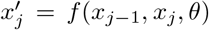 and 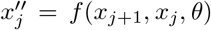, see Fig.1A. The two input pairs having independent and zero means noise, generate two denoised versions of the same *j*^*th*^ image. Then, the final denoised solution 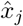 of the *j*^*th*^ image is given by:

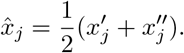

One can view, for every slice *x*_*j*_, the adjacent images *x*_*j*−1_ and *x*_*j*+1_ as discrete versions of the signal along the third direction of the volume. In addition, *x*_*j*−1_ and *x*_*j*+1_ have their own noise that is spatially independent. Due to the FIB-SEM way of imaging (extremely high isotropic resolution along all three dimensions), we consider that the signal between two slices has good enough properties that we can use a Taylor expansion along the third-axis. Thus for every image *x*_*j*_ one can view *x*_*j*−1_ = *s*_*j*_ + *noise* + *e*_0_, where *s*_*j*_ is the real signal, *e*_0_ = *O*(*s*_*j*−1_ − *s*_*j*_), and similarly for *x*_*j*+1_. In such case the eq. 2 can be rewritten as

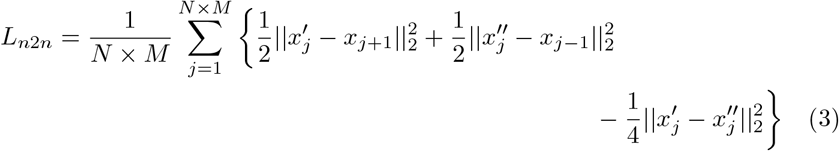

### 2.2 Experiments

The NRRN architecture has been specifically designed for FIB-SEM images denoising, and trained and tested on images acquired with a FEI Helios NanoLab 660 DualBeam using an In-Column Detector (ICD) at Oregon Health & Science University (OHSU) Multi-scale Microscopy Core (MMC). In this section we present the denoising results with the newly proposed NRRN and we also discuss different ways to assess the denoising quality. Furthermore, we compare NRRN outcomes to alternative approaches, we discuss the denoising process and more practical issues like efficiency and transfer learning.

#### 2.2.1 Data and Techniques for Analysis

The OHSU data set is composed of cancer tissue images of dimensions 4*K* × 6*K* pixels with 4*nm* resolution. The way it was acquired allowed the creation of “ground truth” images: the tissue sample surface was scanned 10 times before slicing (the number of scans being limited because of the artifacts produced by the electron beam during scanning), so producing 10 images of the exact same area, and this process was repeated 5 times. The 10 images of the same surface can be considered as technical replicates in which only the noise is different. These images were first registered together using an affine transformation from an in-house stochastic version of TurboReg (Thevenaz, Ruttimann & Unser 1998). Then a ground truth image was generated as follows: for each pixel, the 10 values from the 10 images were sorted, then the 2 most extreme values on each side where removed to avoid outliers, and the remaining 6 values were averaged. The nature of the OHSU data set, with its multiple scans per slice, allows us to examine and compare the way NRRN learns the specific electron microscopy noise and distinguishes it from the true signal.

To analyze the way NRRN deals with volume data and to test on images acquired at a different institution and microscope, we experimented with the publicly available data set from (EPFL Electron Microscopy Dataset n.d.). This data set represents a section taken from the CA1 hippocampal region of the brain, corresponding to a 1065 × 2048 × 1536 pixel volume at 5 × 5 ×5*nm* voxel resolution, where the top 165 slices are used for training and the bottom 165 for validation. Since ground truth image are not available, we employed the data set to examine the efficiency of the proposed algorithm in removing artificially added Gaussian and/or Poisson noise.

To evaluate denoising efficiency, we use two classical measures: the Peak Signal-to-Noise Ratio (PSNR) (Hore & Ziou 2010), and the Structural Similarity Index Measure (SSIM) (Wang, Bovik, Sheikh & Simoncelli 2004). Both measures compare the denoised image to a ground truth image, and in the case of the OHSU data set we are fortunate to have access to acceptable approximations. In addition, we look at the interquartile range (IQR) of the signal across a straight line in an area with flat signal. In the FIB-SEM case, the entire workflow from tissue collection to final image harvesting takes roughly two weeks for 1500 images (Riesterer, López, Stempinski, Williams, Loftis, Stoltz, Thibault, Lanicault, Williams & Gray 2020). During this process the clinical specimen undertakes several epoxy resin infiltration steps to fill the space between the cellular structures. In our analysis we look at the noise presence in the resin because it is a homogeneous material, and as a consequence an “efficient” denoiser should produce a flat signal.

#### 2.2.2 Implementation details, Results and Inference

The OHSU data set was separated into 3 slices for training, 1 for validation and 1 for testing, and the initial large images were cropped into 13650 smaller images of size 256 × 256 pixels, distributed as follow: 8190 images for training (three slices), 2730 for testing and validation (one slice each). The NRRN architecture was implemented in PyTorch, with 4 BUs, the images triplets in the training are taken from the same slice, and the training performed with ADAM optimizer for initial learning rate 10^−4^, *β*1 = 0.9, *β*2 = 0.999, *E* = 10^−8^, and a batch size of 16.

On the test slice, NRRN achieves a PSNR of 31.0197 ± 0.1905 dB and a SSIM of 0.9705 ± 0.0006. Fig. 2 shows an image from the test slice with a PSNR of 31.11dB, significant reduction of the noise on the resin (the IQR is reduced to 0.94 - Fig.2D), while still keeping sharp edges on the cell organelles with reduced noise (see Fig. 2E).

**Figure 2:**
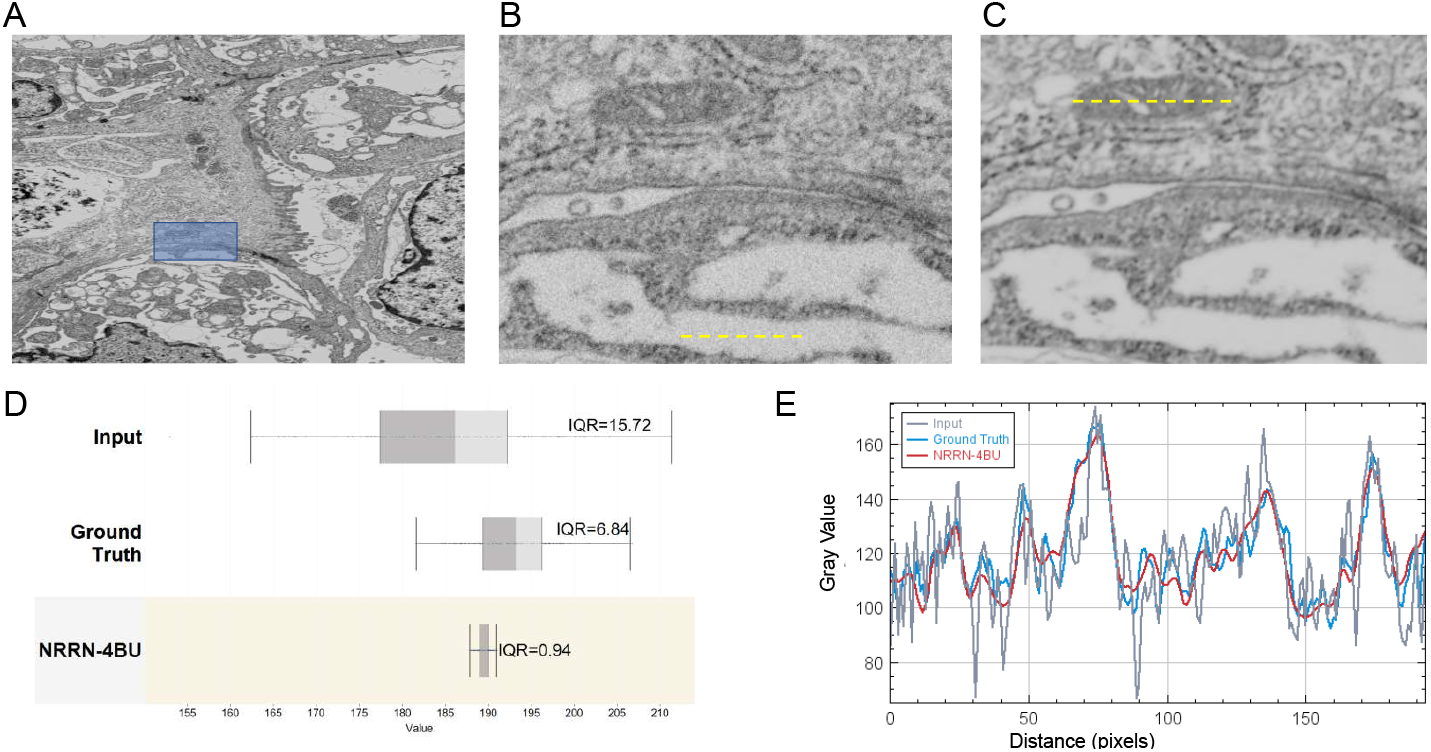
NRRN denoising results, where the yellow doted line shows where image signal is measured. (A) validation image; (B) zoom in of the noisy input image; (C) zoom in of the denoised image; (D) box plot of the signal across the resin (from subfig. B - showing the significant reduction in the signal/noise variation; (E) the signal variation across the mitochondria (from subfig. C)) - the NRRN denoising keeps sharp edges.

Remark: one might argue that using a model taking a triple input is a burden, but double or even single inputs are also possible during inference. However our experiments shows (see Fig. S1, the higher the number of input images, the higher the PSNR and SSIM. Nevertheless, an inference with a double input architecture gives very satisfactory quality results in comparison to the triple input by degrading the overall results (measured in PSNR and SSIM) by only 2%. The signal variation in the flat resin area or across the organelle for two images is almost indistinguishable from the three images. Using a single image or two images would, in addition, reduce the computational burden since we would pass the input through the single branch of the architecture.

#### 2.2.3 Comparisons

We compare NRRN denoising approach on the OHSU data set to the following traditional supervised Noise2Clean training as well as classical training-free methods:

- Training-free methods: BM3D, non-local means (NLM), median and Gaussian filters. We applied the NLM to a single image (here refereed as ‘NLM sngl’) and to an average of 3 images (here refereed as ‘NLM avg’).
- Training methods: U-Net and DenoiseNet. We experimented with the classical DenoiseNet (using ground truth images in the training) as presented in (Remez, Litany, Giryes & Bronstein 2017) (here referred as ‘N2C DenoiseNet-sngl’), but also a version of DenoiseNet that is applicable to a sequence of (three) images (Remez, Litany, Giryes & Bronstein 2018) (here referred as ‘DenoiseNet’ or ‘N2C DenoiseNet’). The U-Net is a classical network with a depth of 4 and batch normalization. Again, we trained on a single image and on an average of three consecutive images (as input) and the corresponding ground truth image - here referred as ‘N2C U-Net sngl’ and ‘N2C U-Net avg’ (or simply U-Net) respectively.

The OHSU FIB-SEM input images of size 4*K* × 6*K* are rather large to be fit into the GPU memory. Since the noise is random at the pixel level (no spatial dependencies), during inference we crop the large images into 345 smaller patches of size 256 × 256, denoise them and stitch them back together to reconstruct the large image, allowing a 20 pixels overlap to avoid the edge effect introduced by convolution layers. In contrast to (Wu, Silversmith & Seung 2019) where authors weighted the overlapping values, we cut them out. This does result in additional patches that need to be denoised, but it overcomes border artifacts. On Fig. 3A, we plotted the patches PSNR distribution for each of the compared architectures. One advantage of NRRN is that the noise is removed homogeneously across all patches, with a consistent PSNR of 31.21 ± 0.52, which is marginally better than DenoiseNet 0.58 standard deviation, but significantly outperforms the U-Net deviation of 1.26.

**Figure 3:**
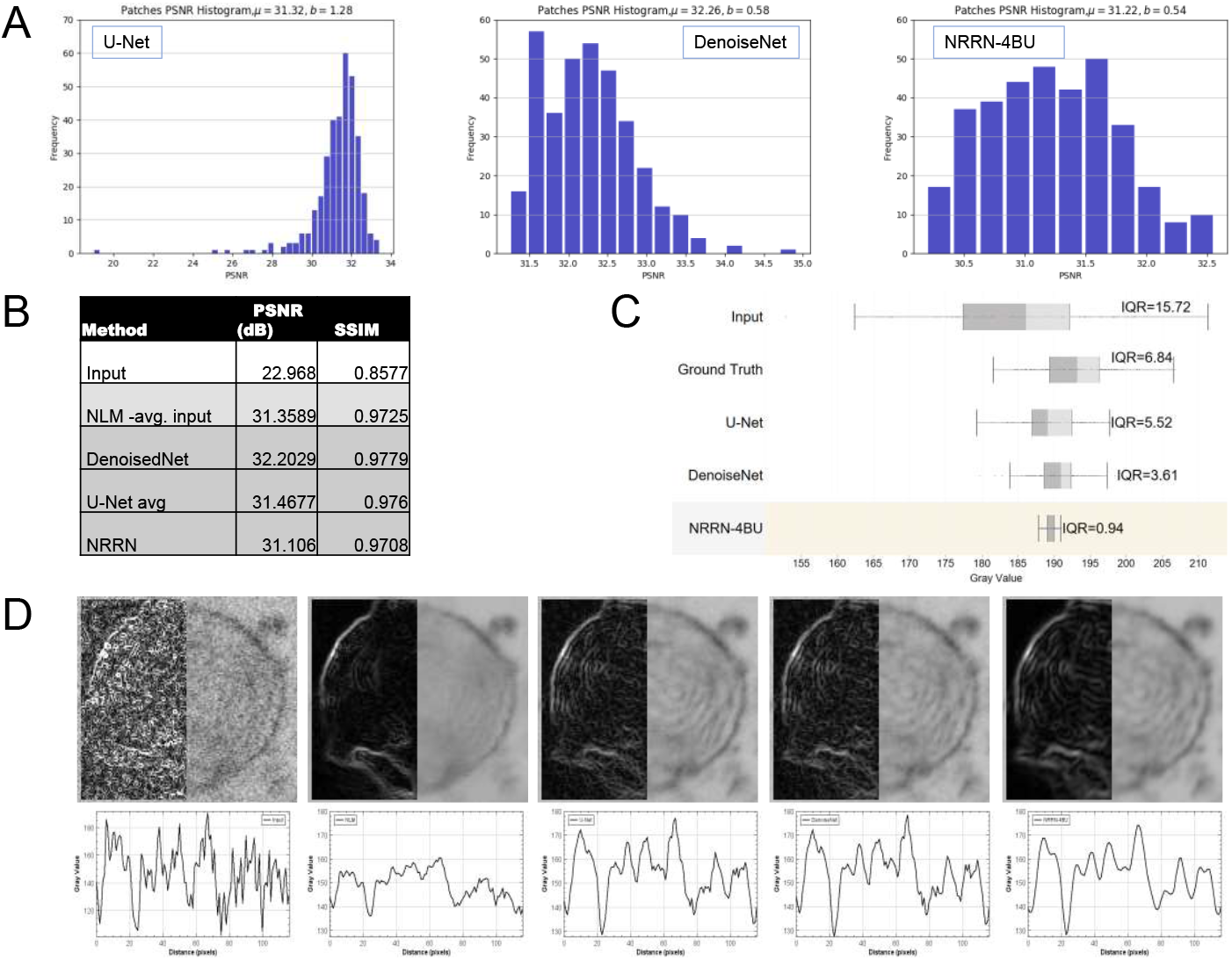
Comparison of NRRN to other alternative denoisers. (A) PSNR Histogram for the patches constructing a FIB-EM image, from left to right U-Net, DenoiseNet, NRRN; (B) Comparison of the PSNR and SSIM; (C) Comparison of the signal across the resin for U-Net, DenoiseNet, NRRN and the Ground Truth image. The IQR of NRRN outperform the rest of the networks; (D) Variation of the signal across a cell organelle and edge sharpness: Input image, NLM, U-Net, DenoiseNet and NRRN (from left to right).

In this paper we argue that the loss function as defined in eq. (3) results in better denoising. The ground truth images have better signal than the input images, but they are not noise free. Using per-pixel loss with a ground truth in the training (in the face of L1 or L2 norm) would maximize the PSNR but results in an image that inherits the noise present in the “ground truth”. This is not the case of NRRN with the loss function defined as in eq.(3), we are able to better remove the random pixel noise. On Fig. 3C, we compared the noise in the signal flat resin region for U-Net, DenoiseNet and NRRN. Even though DenoiseNet and U-Net significant improvement in comparison to the input image and the ground truth, NRRN significantly outperform them all.

Comparisons on the final whole image based on PSNR and SSIM are shown in table 1. The non-local mean (NLM-avg), DenoiseNet and U-Net all outperform NRRN, in terms of PSNR and SSIM. That is expected since DenoiseNet and U-Net were trained to minimize the MSE loss (equivalent to maximizing the PSNR). In the case of non-local mean the high PSNR is achieved by aggressively flattening the signal even on biological structures of interest. NRRN does a better job in removing noise across the resin while sharpening the biological structures - visible on Fig. 3C-D.

**Table 1:** Denoising performance of NRRN vs other denoisers in terms of PSNR and SSIM on the OHSU data set. The comparison is carried out for BM3D, Non-Local Means Denoising (NLM) for a single and sequence of 3 images (NLM - avg input), and Gaussian Filter, Median Filter, U-Net DenoiseNet and NRRN (the best results are highlighted in bold font).

| Method | PSNR<br>(dB) | SSIM |
| --- | --- | --- |
| Input | 22.9680 | 0.8577 |
| NLM -avg. input | **31.3589** | **0.9725** |
| BM3D | 28.2515 | 0.8747 |
| Median Filter-avg. input | 28.0214 | 0.9495 |
| Gauss Filter-avg. input | 27.0649 | 0.9195 |
| NLM -sngl | 24.0703 | 0.6195 |
| N2C DenoiseNet | **32.2029** | **0.9779** |
| N2C U-Net avg | 31.4677 | 0.9760 |
| *NRRN* | 31.1060 | 0.9708 |
| NRRN -2 img | 30.4194 | 0.9666 |
| N2C U-Net sngl | 29.1844 | 0.9571 |
| N2C DenoiseNet -sngl | 28.3436 | 0.9559 |

Besides the OHSU data, we synthesized additional noise to the EPFL data set. We fine tuned the OHSU pre-trained models by continuing the training on the small EPFL volume data set. During training, we corrupt the EPFL images with Poisson noise for random peak values ranging in [1, 50] and also added Gaussian noise with random *σ* between 10 and 75. The results of denoising images from the validation data set with added Poisson and Gaussian noise respectively are in Tables 4A and 4B. In terms of the quantitative measure PSNR, NRRN with its unsupervised training shows very comparable performance to the supervised training - marginal improvement in some of the cases and a slight under-performance in others. A closer visual inspection (see Fig. 4C) reveals that: U-Net tends to over smooth the details; DenoiseNet adds white speckles for low peak values and high *σ*, indicating that the network struggles to extract enough information from the significantly damaged images; while NRRN does not display any of these issues because the Building Units are able to synthesize the information from the adjacent slices and reconstruct and remove the noise without over smoothing or adding artifacts.

**Figure 4:**
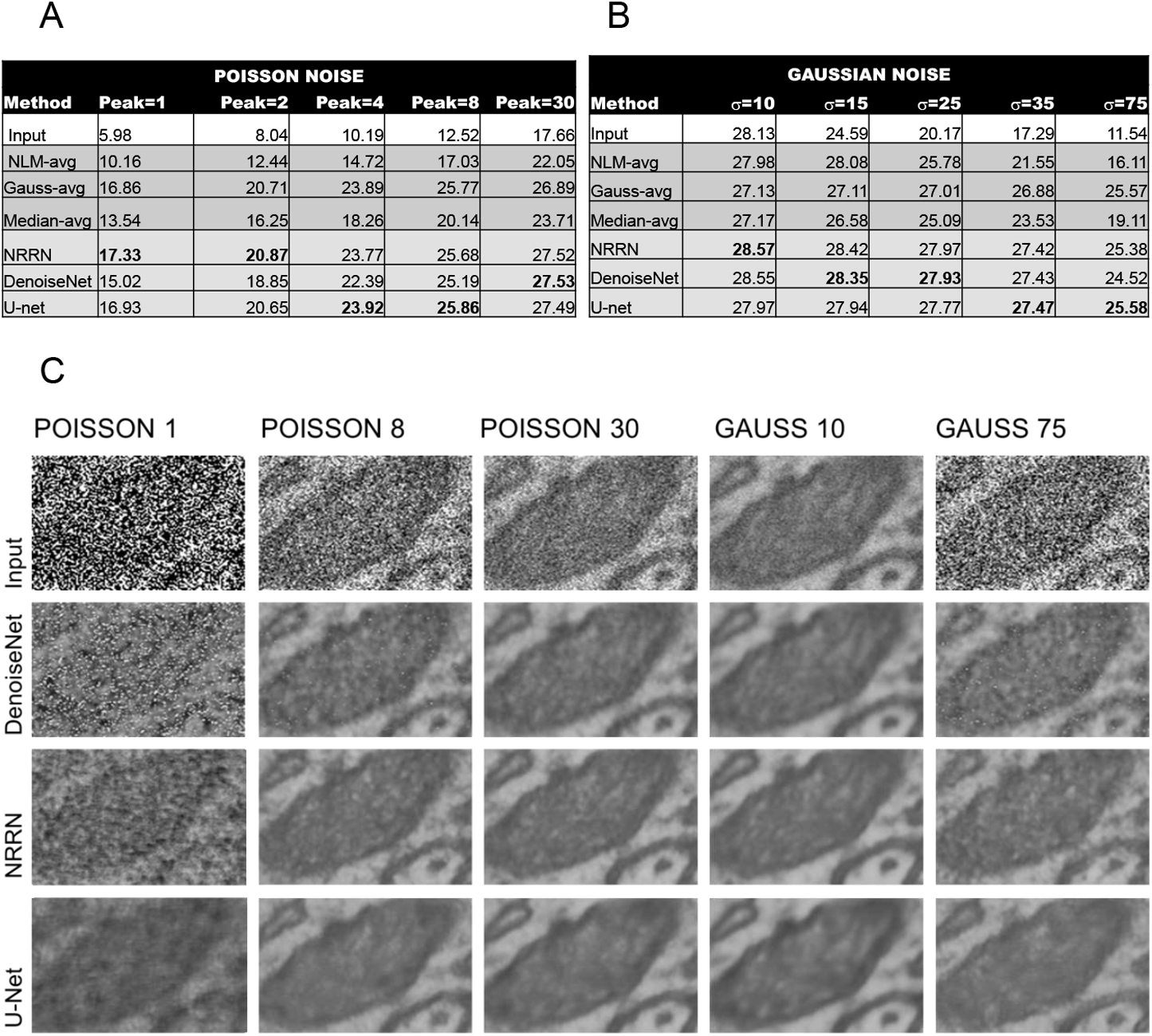
Denoising EPFL images. (A) PSNR in dB for EPFL image with added Poisson noise with peak=1, 2, 4, 8, 30; (B) PSNR in dB for EPFL image with added Gaussian noise with *σ* = 10, 15, 25, 35, 75; (C) Comparison of the denoising performance. Top to bottom: Input image, DenoiseNet, NRRN and U-Net. Left to right: added Poisson noise with *peak* = 1, 8, 30 and Gaussian noise with *σ* = 10, 75.

#### 2.2.4 Exploring the denoising process

Similar to DenoiseNet, our algorithm allows intermediate noise estimates after each Building Unit. Our exploration of the denoising process is carried out for NRRN with 5 BUs on both data sets: OHSU and EPFL. Each BU SSIM shows that the majority of the denoising happens in the first 1-3 layers where we observe a more significant and gradual improvement (see Fig. 5A, when the last two BUs become important for highly corrupted images (Poisson noise with peak=1 and Gaussian noise *σ* = 75). On Fig. 5B we looked at the signal across the resin. We scaled the signal with the signal mean value and plotted the scaled input image signal (x-axis) versus the scaled BUs signal (y-axis). We see that the signal (which on the resin area is mostly noise) gradually fattens.

**Figure 5:**
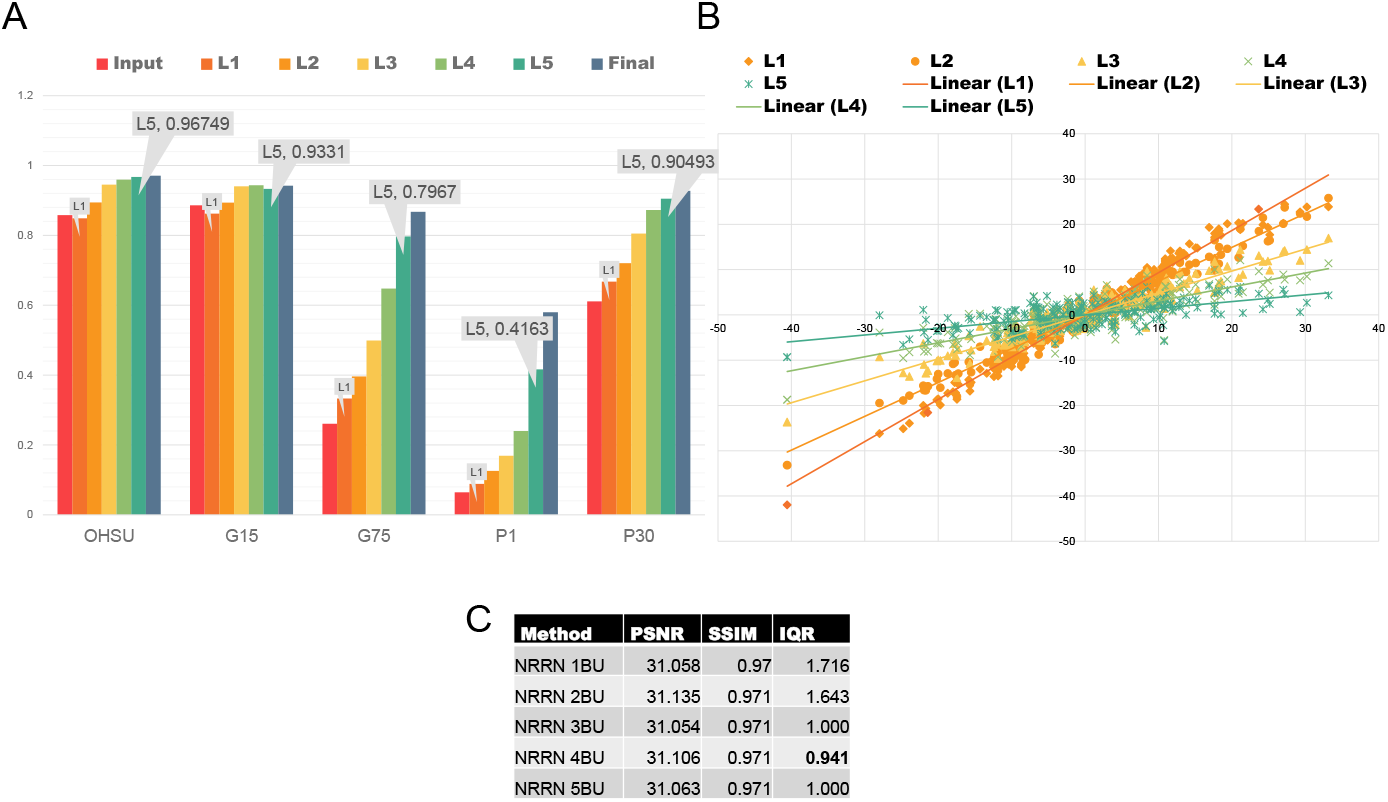
Iterative denoising process. (A) SSIM at each layer for OHSU data set and EPFL with added Poisson noise with peak 1 and 30 (P1, P30) and added Gaussian Noise with *σ* = 15, 75 (G15,G75); (B) Input signal scatter plot across the resin vs layers signal; (C) PSNR, SSIM and IQR for the NRRN family

The ability to look at the intermediate denoising results and the modular architecture helps with the network fine tuning: the higher the level of noise, the higher the number of BUs. To investigate such behavior, we trained a whole family of NRRN networks ranging the number of BUs from 1 to 5 on the OHSU data set. We measured the signal variation on the resin in terms of IQR, PSNR, and SSIM. Table 5C indicates that while the SSIM and PSNR seems to not improve with more BUs, the signal on the resin flattens (smaller IQR) until the 4^*th*^ BU.

#### 2.2.5 Transfer learning and efficiency

In the case of biomedical images the data sets available for training are rather small. Transfer learning with pre-training on a bigger data set works well: to denoise the EPFL data set, we pre-trained on the OHSU data set and then fine-tuned it on a small portion made of 165 images of the EPFL data set. We also experimented with directly training on the EPFL data set. The NRRN networks (even with just a single BU) are significantly better in extracting the information of interest than DenoiseNet and U-Net - see Table 2 (as we have seen in Tables 4A-B the three architectures achieve similar results on bigger data set).

**Table 2:** Denoiser performance without transfer learning on the small EPFL data set. Quantification in terms of PSNR (in dB) shows that NRRN outperforms the other networks (the best results are highlighted in bold font).

| Method | Poisson<br>peak=1 | Poisson<br>peak=30 | Gaussian<br>$\sigma = 75$ | Gaussian<br>$\sigma = 10$ |
| --- | --- | --- | --- | --- |
| NRRN 1BU | **17.8976** | **27.5081** | **25.1841** | **28.4803** |
| DenoiseNet | 9.4712 | 19.6979 | 14.3589 | 25.2942 |
| U-Net | 9.4189 | 20.9759 | 15.0638 | 27.1999 |

Furthermore, we studied the giga operations per second (GOPs) and number of parameters vs one of the quality measures for the NRRN family (single branch), (N2C) U-Net and (N2C) DenoiseNet (see Fig. S2). If we are to find the best trade off denoising quality versus efficiency, then the NRRN with 3 Building Units gives, in our opinion, the best trade off from the whole NRRN family, DenoiseNet, and U-Net for the OHSU data set. For more noise corrupted images (as it was for EPFL with Poisson noise with peak 1-8 or Gaussian Noise with *σ* = 75) the 5 BUs with their bigger receptive field would be required.

### 2.3 Denoising advantages and new possibilities

Thanks to the noise2noise approach adopted by the NRRN architecture, one can train a denoising model that would be specific to each data set. Such possibility provides a solution to efficiently denoise every data set in a unique and dedicated manner. The efficiency reached by NRRN highly reduces the noise while preserving edges and improving the contrast of shallow structures of interest like mitochondria texture or macropinosomes (see figure 3 D). Such improvements allow for a better visualization and as a consequence understanding of biological structures, as well as an easier manual or automatic segmentation. Moreover, denoising improves texture characterization and then classification, for example the nuclei chromatin, by reducing the texture features instability which is often introduced by noise and artifacts as it is the case for statistical matrices like Haralick features or size zone matrix (Thibault & Shafran 2016).

## 3 Discussion

The advances in the focused ion beam-scanning electron microscopy provide high resolution cell images, which unfortunately contain a significant quantity of noise. In this paper, we discussed the advantage and further developed the idea of using a sequence of images with a noise2noise approach for removing the noise. This approach has the benefit of not requiring ground truth images for training, which is typically the case for biomedical deep learning tackled problems. We suggested a novel modular architecture (NRRN) that is able to exploits the sequential nature of the images to reconstructs and remove the noise from the original image. We demonstrated that this unsupervised approach leads to consistent noise removal across the entire image without regard to the structures present. Besides, we showed that the architecture allows a glimpse into the denoising process that can be used to adjust the depth of the network to the available computational resources and/or presence of noise.

## 4 Acknowledgments

FIB-SEM data included in this manuscript was generated at the Multiscale Microscopy Core (MMC), an OHSU University Shared Resource, with technical support from the OHSU Center for Spatial Systems Biomedicine (OCSSB). Specimen acquisition support from the SMMART clinical coordination team was invaluable. This manuscript was supported by Prospect Creek Foundation, the Brenden-Colson Center for Pancreatic Care, the NCI Cancer Systems Biology Measuring, Modeling, and Controlling Heterogeneity (M2CH) Center Grant (5U54CA209988), the NCI Human Tumor Atlas Network (HTAN) Omic and Multidimensional Spatial (OMS) Atlas Center Grant (5U2CCA233280), the OHSU Knight Cancer Institute NCI Cancer Center Support Grant (P30CA069533), and the OCSSB.

This study was approved by the Oregon Health & Science University Institutional Review Board (IRB # 16113). Participant eligibility was determined by the enrolling physician and informed consent was obtained prior to all study protocol related procedures.

We thank Eugenio Culurciello and the rest of the Micron ML team for their useful comments and discussions.

## 5 Author Contributions

K.G., A.Ch., B.DeLaR. designed the architecture; J.R. and E.S. prepared the tissue samples and collected the data images; K.G., A.Ch., B.DeLaR., G.Th., T.Lo designed the experiments and analyzed the results; J.W.G. conceived the project. All authors have read, edited, and approved the content of the manuscript.

## Supplemental Figures

**Figure S1:**
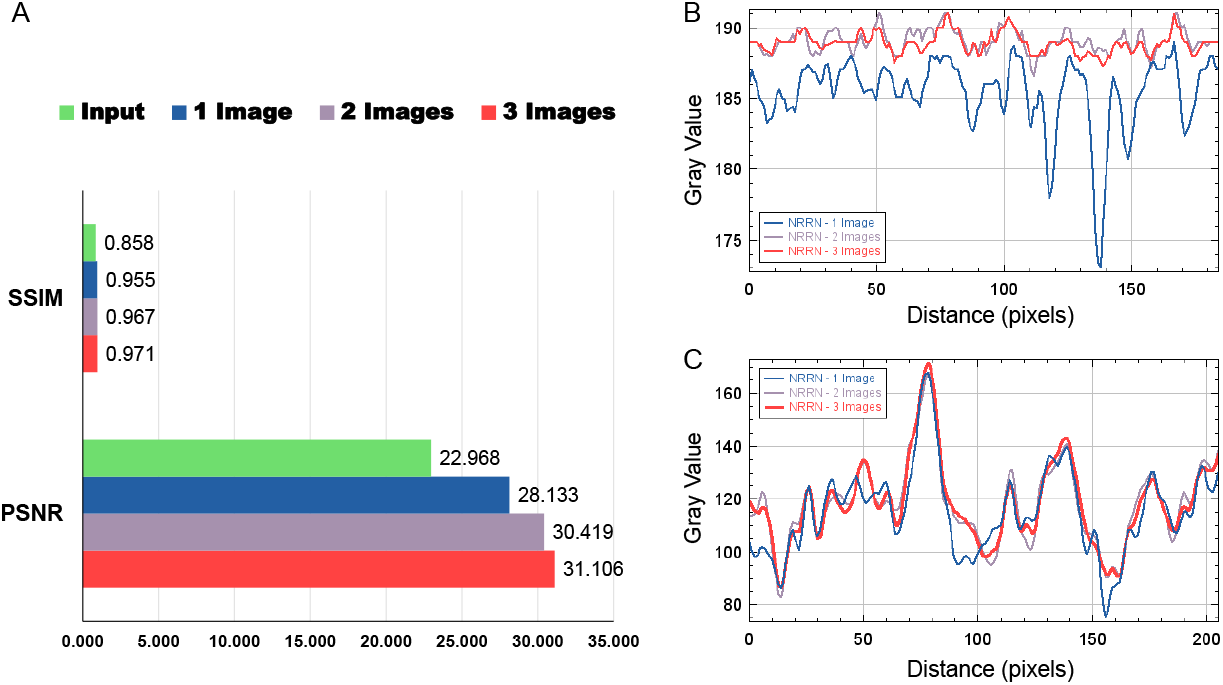
Related to Section 2.2.2 and the remark on inference with less than three images. Quantitative analysis for inference with 1, 2 and 3 input images. (A) PSNR and SSIM for inference with 1, 2 and 3 input images. (B) The signal across the resin for inference with one, two and three images. (C) The signal across an organelle.

### Peak Signal to Noise Ratio - PSNR

PSNR measures the pixel distance between two images. Given a reference image *f* and a test image *g*, both of size *M* × *N*, the PSNR between *f* and *g* is defined by:

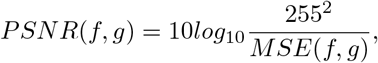

Where

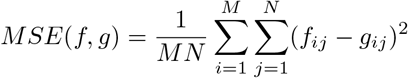

### Structural Similarity Index Measure - SSIM

SSIM measures the perceptional difference between two images. Let’s consider a reference image *f* and a test image *g*, both of size *M* × *N* and denote with *µ*_*f*_ the mean value of the images *f*, *σ*_*f*_ the standard deviation of f, *σ*_*fg*_ the covariance between *f* and *g*. The SSIM between *f* and *g* is defined by:

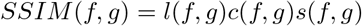

where

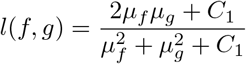

is the luminance function which compares the closeness of the two images’ mean luminance (*µ*_*f*_ and *µ*_*g*_);

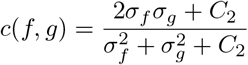

is the contrast comparison function which measures the closeness of the contrast (*σ*_*f*_ and *σ*_*g*_) of the two images;

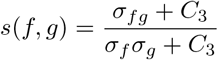

is the structure comparison function which measures the correlation coefficient (*σ*_*fg*_) between the two images f and g.

**Figure S2:**
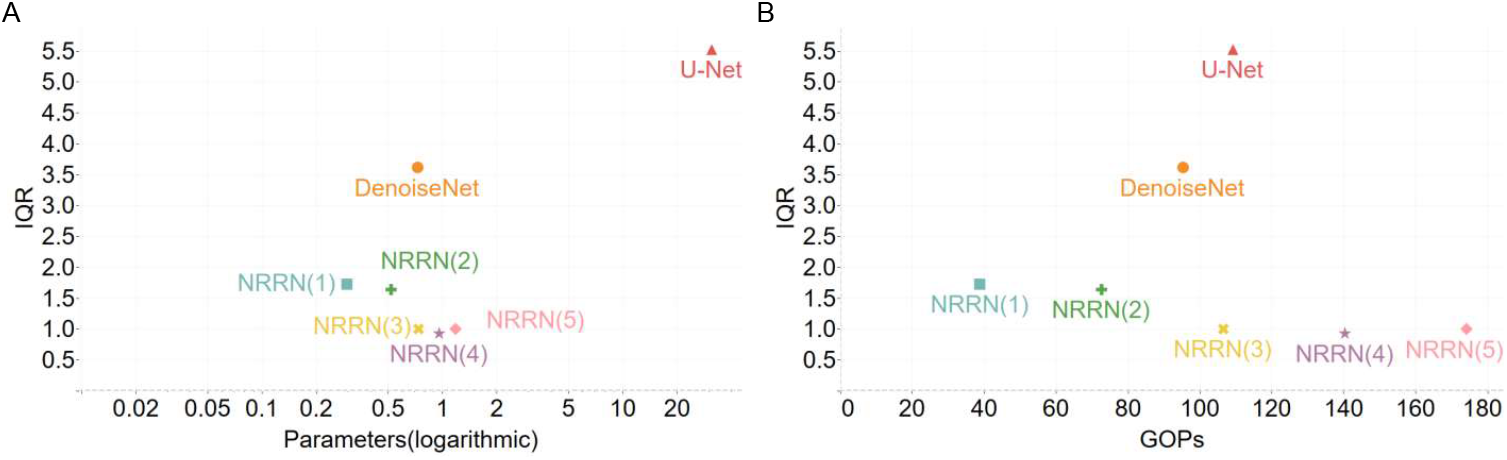
Related to Section 2.2.5 and more specifically the efficiency of the NRRN. Comparison of the GOPs and number of Parameters (Million) vs IQR for NRRN family, DenoiseNet, and U-Net. (A) Number of parameters vs. IQR across the resin. (B) Number of giga operations (GOPs) vs. IQR. For the NRRN Family the GOPs are for only one (of the two) branches, the noise2noise training requires two branches which will double the GOPs but avoids the need of ground truth images.

SSIM takes values between 0 and 1, where a value of 0 means no correlation between the two images, and 1 means that *f* = *g*.

### Rectified Linear Unit - ReLU

ReLU is a non-linear activation function that is used in multi-layer neural networks or deep neural networks (Glorot, Bordes & Bengio 2011). This function is defined as:

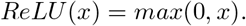

